# Highly effective modulator therapy hinders the emergence of hyperbiofilm variants of *Pseudomonas aeruginosa* PA14 grown in a cystic fibrosis lung model

**DOI:** 10.64898/2026.08.08.743698

**Authors:** Émile Létourneau, Océane Goncalves, Jean-Philippe Côté, Fabrice Jean-Pierre

**Author notes:** Corresponding authors (J-P.C), (F.J-P). These authors contributed equally to this work (Émile Létourneau, Océane Goncalves).

## Abstract

*Pseudomonas aeruginosa* is an opportunistic pathogen that often adopts persistent phenotypes — such as biofilm formation— that are associated with chronic infections including those observed in the cystic fibrosis (CF) lung Recently, highly effective modulator therapy (HEMT) such as elexacaftor/tezacaftor/ivacaftor (ETI) has significantly improved the quality of life of people with CF (pwCF). Yet a potential direct impact of ETI on the physiology of *P. aeruginosa* during growth to a remodeled CF lung environment has remained unexplored. To address this, we conducted an experimental evolution using *P. aeruginosa* PA14 grown in CF-like conditions in the presence or absence of ETI. We observed a marked reduction in biofilm formation and in the number of small colony variants (SCVs) for *P. aeruginosa* populations evolved under ETI treatment. Also, sequencing of specific evolved clones exhibiting distinct morphotypes revealed two major observations: (i) *P. aeruginosa*-evolved communities exposed to ETI retained a wild type-like morphotype and, (ii) *P. aeruginosa* populations evolved in the absence of ETI adopted a SCV-like phenotype with mutations acquired in the Wsp chemosensory pathway. Furthermore, analysis of evolved populations revealed that ETI treatment likely modulates c-di-GMP pools by driving mutations in an enzyme catalyzing the degradation of this second messenger. Overall, our work suggests that ETI has the potential to hinder the acute to chronic biofilm transition of *P. aeruginosa* thereby limiting the emergence of variants typically associated with long-term CF lung colonization.

## Introduction

Cystic fibrosis (CF) is a genetic disease caused by mutations in the CF transmembrane conductance regulator (CFTR), resulting in the accumulation of a thick mucus in the lungs of people with CF (pwCF) [1]. This modified airway environment promotes chronic colonization by pathogens –such as *Pseudomonas aeruginosa*– which persists through the formation of biofilm-like communities. Furthermore, while no treatment strategies thus far have resulted in the complete eradication of *P. aeruginosa* from the CF lung, several groups have reported the emergence of genetic variation in *P. aeruginosa* populations that colonize the CF airways over many years [2,3]. Therefore, these observations strongly suggest that *P. aeruginosa* can respond to its environment by adapting to its host during an infection, thereby contributing to worsened patient outcomes [4].

Recently, highly effective CFTR modulator therapy (HEMT) such as elexacaftor/tezacaftor/ivacaftor (ETI), has been developed to partially restore malfunctioning CFTR activity, the underlying cause of the disease in individuals carrying an eligible mutation [5]. This resulted in significantly improved outcomes in lung function and quality of life of pwCF [6]. Interestingly, although ETI therapy substantially reduces the total bacterial burden and the relative abundance of *P. aeruginosa* in the airways of pwCF, chronic infections caused by this pathogen persist after multiple years of treatment [7,8]. Furthermore, most patients remain infected across multiple lung segments, with the same clonal lineages continuing to evolve within the ETI-treated respiratory environment [9,10]. Therefore, these multiple lines of evidence highlight the need to investigate long-term effects of CFTR modulators on *P. aeruginosa* persistence and adaptation in the CF airway.

To address this knowledge gap, we used a reductionist approach. That is, we exposed *P. aeruginosa* to ETI therapy over the course of 20 days using CF-like conditions and looked at the potential impact of this CFTR-specific treatment on biofilm formation, a critical pathogenesis component in *P. aeruginosa*. We observed that *P. aeruginosa* populations exposed to ETI treatment are hindered in their capacity to generate hyperbiofilm-forming variants that are typically associated with long-term CF lung colonization.

## Results and Discussion

### ETI represses the emergence of hyperbiofilm-forming variants of *P. aeruginosa*

*P. aeruginosa* wild-type (WT) strain PA14 grown in artificial sputum medium (ASM) was exposed to physiologically relevant concentrations of ETI for 20 days (with passages every 48 h) and compared their evolution to DMSO-treated control biofilms. At each passage, cultures were plated onto LB agar to assess colony morphology, and each evolved populations were preserved as glycerol stocks for subsequent analysis. After 5 passages, we observed a marked increase in biofilm formation in 5/10 DMSO-treated control lineages (Fig 1A and 1B). In contrast, none of the ETI-treated lineages showed a similar increase in biofilm formation (Fig 1A and 1B). Throughout the evolution, we detected colonies with distinct morphotypes, many of which appeared significantly smaller than WT PA14 (Fig 1C). These smaller colonies displayed characteristics that are reminiscent of small colony variants (SCVs), including a rugose morphology with enhanced Congo Red binding and absence of motility (Fig 1D). Interestingly, increased biofilm formation correlated with SCVs frequency in evolved lineages, where the high-biofilm DMSO-treated control lineages contained high frequencies of SCVs (Fig 1E). Furthermore, ETI-exposed lineages exhibited an almost complete absence of SCV (Fig 1E). Taken together, these initial observations support a model where ETI treatment impacts morphological heterogeneity and biofilm formation of *P. aeruginosa*.

**Fig 1.**
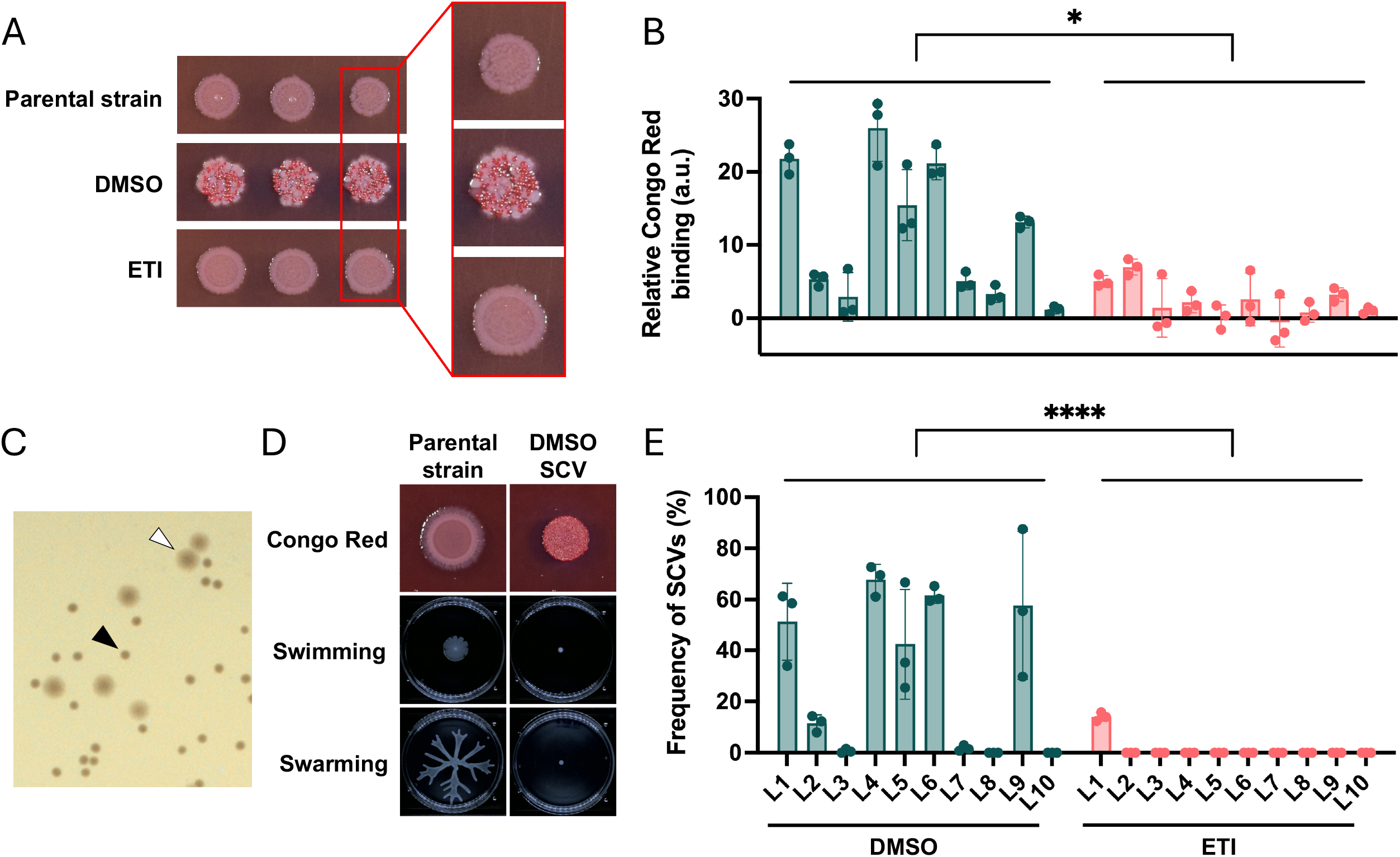
Impact of ETI on biofilm formation and small colony variant emergence in *P. aeruginosa* PA14 evolved lineages. (**A**) Representative images from the Congo Red binding assay for the *P. aeruginosa* PA14 parental strain and evolved strains of lineage 6 (L6) after passage 5. (**B**) Quantification of Congo Red binding for all 20 lineages after passage 5, under both DMSO (dark green) and ETI (pink) conditions, compared to the parental strain. Biofilms of each lineage were tested in triplicate. Statistical analysis was performed using a Welch Two Sample t-test to compare mean CR binding value between DMSO-treated control lineages and ETI-treated lineages; \**p* ≤ 0.05. (**C**) L6 population was recovered from glycerol stocks. Black arrow indicates SCV, while white arrow indicates parental-like colony. (**D**) Representative images of SCV Congo Red binding, swarming, and swimming motility compared to the PA14 parental strain. (**E**) Quantification of SCVs frequency across lineages after passage 5. The experiment was performed in triplicate. Statistical analysis was conducted using a Wilcoxon rank-sum test with continuity correction; \*\*\*\**p* ≤ 0.0001.

### ETI impacts biofilm formation through changes in c-di-GMP pools

To further characterize the molecular basis of these divergent adaptive outcomes, we sequenced single evolved clones of *P. aeruginosa* exhibiting distinct morphotypes. These clones were selected from the three lineages displaying the most pronounced Congo red staining difference (Fig 1B – L4, L6, and L9 at passage 5). For the DMSO-treated control lineages, we isolated and sequenced three independent clones exhibiting a parental-like morphotype and three independent SCV clones. For comparison, we also isolated and sequenced three independent clones displaying a parental-like morphotype from ETI-treated lineages. Of note, these ETI-treated lineages did not contain SCVs. Differences from the parental strain were identified using two complementary variant-calling pipelines (Medaka and Clair3) to identify SNPs, insertions and deletions (Fig 2A and S1 Data). Our analyses revealed two mutually exclusive mutational patterns. First, in the DMSO-treated control SCV clones, mutations were consistently mapped to the Wsp chemosensory pathway across all three independent sequenced lineages. More specifically, *wspE* harbored a Lys748Asn substitution in all three SCV clones from L4, while *wspC* and *wspD* each carried missense mutations in all SCV clones from L6 and L9 (Fig 2A and S1 Data). The Wsp pathway controls WspR-dependent c-di-GMP synthesis upon surface contact, and gain-of-function mutations in this system are well-established as drivers of the SCV and hyperbiofilm phenotypes in *P. aeruginosa*, including clinical isolates of pwCF [11,12]. In contrast, ETI-treated parental-like clones harbored mutations in *dipA*, which encodes a c-di-GMP phosphodiesterase involved in biofilm dispersion in *P. aeruginosa* [13]. All three ETI-treated clones from L6 contained a *dipA* Asp796Glu mutation (S1 Data). One clone from L4 contained a *dipA* Asp269Ala mutation, and we identified a *dipA* frameshift (Ala13fs) in two of three clones from L9 (Fig 2A, S1 Data). Of note, no *wsp* mutations were identified in ETI-treated clones, while no mutations in *dipA* were observed in DMSO-treated control SCV clones (S1 Data). This suggests that ETI exposure is likely driving a separate adaptive trajectory, distinct from the Wsp-mediated SCV pathway observed in DMSO-treated control lineages. Interestingly, *dipA* variants were also detected in parental-like clones 1 and 3 from DMSO-treated control lineage 6 (S1 Data). Together, our results suggest that both adaptive responses may co-emerge within evolving populations, but that ETI treatment hinders the emergence of mutations in the Wsp system which is known to drive hyperbiofilm formation in *P. aeruginosa* (Fig 2A).

**Fig 2.**
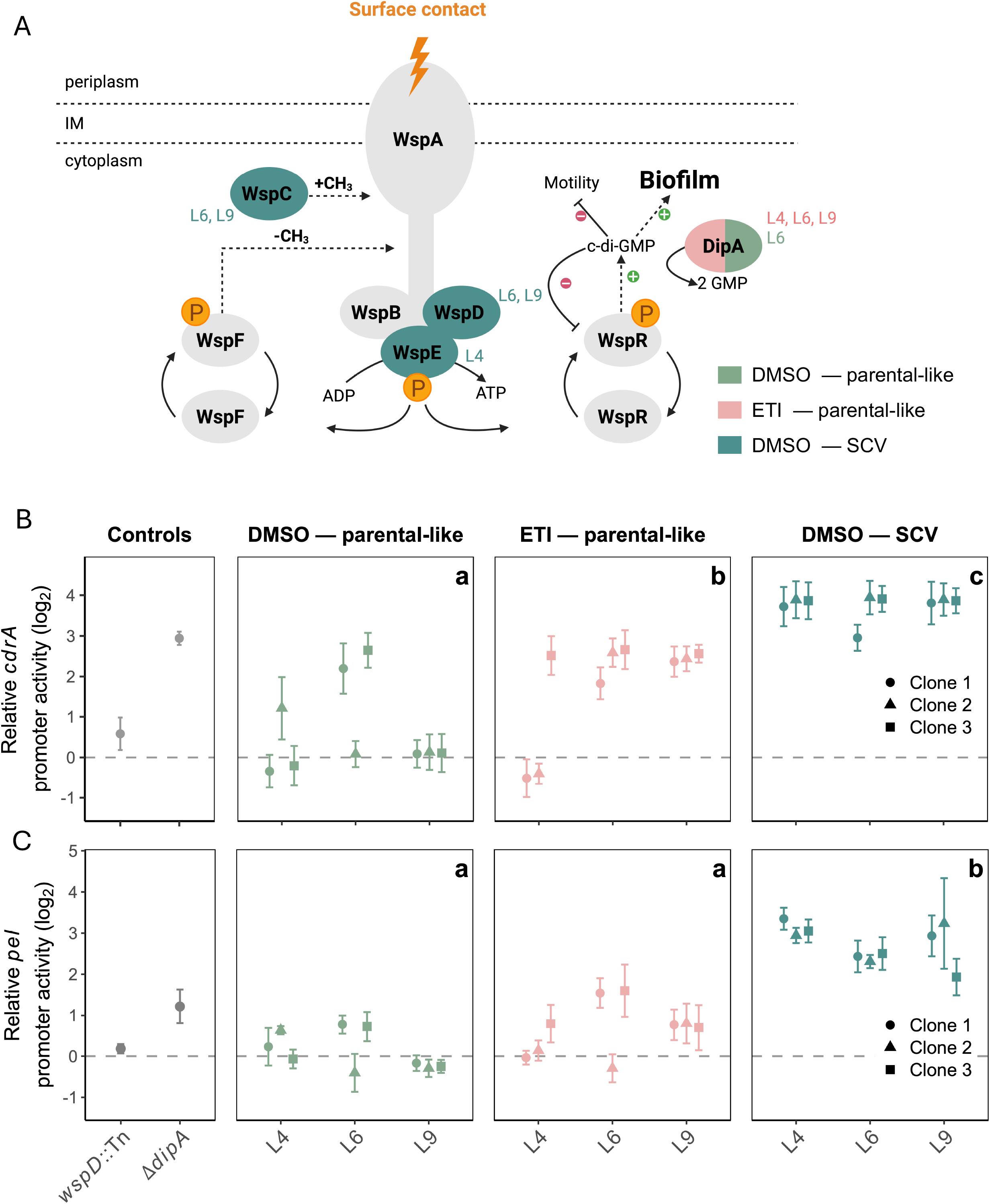
Distinct mutational routes to altered c-di-GMP signalling in DMSO-treated control and ETI-treated *P. aeruginosa* lineages. (**A**) Schematic representation of the Wsp chemosensory pathway and DipA-mediated cdi-GMP regulation in *P. aeruginosa*. Colored ellipses indicate mutations identified by whole-genome sequencing of individual clones isolated at passage 5 from lineages L4, L6, and L9: dark green, mutations in SCV clones from DMSO-treated control lineages (*wspC, wspD, wspE*); pink, mutations in ETI-treated parental-like clones (*dipA*); green, mutations in parental-like clones from DMSO-treated control lineages (*dipA*). See S1 Table for complete variant details. (**B & C**) Relative *cdrA* and *pel* promoter activity respectively in individual evolved clones isolated at passage 5 from lineages L4, L6, and L9. Luminescence and fluorescence are expressed as the log2 of the area under the curve (AUC) ratio relative to the parental PA14 strain. Each point represents the mean of three biological replicates for one independent clone; error bars indicate ± SD. Colors match those of Figure 1A, and grey ellipses represent the controls. The dashed line at y = 0 indicates the level of the parental PA14 strain. Statistical differences between groups were assessed using a linear mixed-effects model followed by post hoc Tukey tests. Letters above data points indicate statistically distinct groups (*p* < 0.05).

To further assess whether these mutational profiles resulted into differential c-di-GMP signalling and biofilm production, we introduced *cdrA*::*lux* and *pel*-GFP transcriptional reporters into each evolved clone. Then, we quantified *cdrA* promoter activity as a validated proxy for intracellular c-di-GMP levels and GFP fluorescence as an indirect readout of Pel exopolysaccharide production [14,15]. Both reporters yielded concordant results (Fig 2B and 2C). That is, SCV clones displayed elevated *cdrA* and *pel* promoter activity– approximately 8 –16 fold above the parental PA14 strain– consistent with constitutive Wsp pathway activation and sustained c-di-GMP accumulation. Parental-like clones from DMSO-evolved lineages showed *cdrA* and *pel* promoter activity comparable to the parental PA14 strain (Fig 2B and 2C). However, ETI-treated parental-like clones harboring *dipA* variants displayed intermediate promoter activity for both reporters –higher than DMSO controls but lower than SCV clones– consistent with the moderate c-di-GMP elevation expected from loss of DipA phosphodiesterase activity [13]. ETI-treated clones lacking *dipA* mutations, as well as the rare *dipA*-mutant clones identified in DMSO control lineages, mirrored this pattern, reinforcing the genotype–phenotype link (Fig 2B and 2C). Of note, clone 2 from L6 is an exception as low level *pel*-GFP expression was observed despite displaying elevated c-di-GMP production (Fig 2C). Together, these findings indicate that two adaptive trajectories toward increased c-di-GMP and biofilm formation emerged during experimental evolution. First, in the absence of ETI the Wsp pathway is preferentially targeted, driving SCV emergence with high c-di-GMP and a marked hyperbiofilm phenotype. Second, in the presence of ETI this trajectory is constrained and *dipA* variants are likely instead selected, producing a moderate c-di-GMP elevation insufficient to drive full *pel-*mediated hyperbiofilm formation and SCV conversion [16,17].

Collectively, our findings suggest that ETI exposure alters the adaptive landscape of *P. aeruginosa* during evolution in a CF-like environment. We report that ETI exposure limits the emergence of Wsp-driven SCV and hyperbiofilm-forming variants while still permitting an alternative adaptive trajectory through the generation of *dipA* variants which are associated with moderate c-di-GMP signalling. These results suggest that HEMT may directly influence the long-term evolutionary dynamics of *P. aeruginosa* in the CF airway and limit its capacity to adopt recalcitrant phenotypes such as SCVs. Future work will include whole-population sequencing of each evolutionary lineage to capture the full spectrum of genetic diversity and to determine whether the mutational patterns identified at the clonal level reflect broader adaptive trends across the evolving populations. Together, our work provides an important experimental framework for understanding how HEMT reshapes bacterial adaptation and highlights c-di-GMP signalling as a key axis through which *P. aeruginosa* navigates the remodelled CF lung environment.

## Materials and methods

### Strains and growth conditions

*Pseudomonas aeruginosa* PA14 (FJP1) was used for all experiments [18]. Bacterial strains were cultured in Artificial Sputum Medium (ASM) as previously described [19]. The CFTR modulators combination elexacaftor/tezacaftor/ivacaftor (ETI, final concentration of 3 µM, 18 µM and 1 µM for each component) was dissolved in dimethyl sulfoxide (DMSO) to prepare a 1000x stock solution. During the adaptive evolution experiment, ETI was added to the cultures at a final 1x concentration. A clean *dipA* deletion mutant of PA14 (FJP432) was used as a control in reporter assays. The p*cdrA*::*lux* reporter plasmid (pMS402) was maintained under kanamycin selection (150 µg/mL).

### Experimental evolution assay

Experimental evolution of *P. aeruginosa* was conducted over 20 days using 96-well plates under aerobic conditions at 37 °C. Twenty independent evolutionary lineages were treated with ETI or a DMSO control. Bacterial cultures were passaged every 48 hours. Supernatants were removed, and the biofilms were washed twice with PBS and resuspended in 100 µL of ASM 1x. 1 µL of the resuspended biofilms (corresponding to approximately 10^5^ cells) was transferred into 99 µL of fresh ASM into a new 96-well plate. In parallel, at each passage, 30 µL of a 10^-5^ and 10^-7^ dilution of the resuspended biofilms were plated on LB agar and colony morphology was assessed after 18 hours of incubation at 37 °C. Distinct morphotypes were isolated and preserved in addition to total populations. Frozen stocks (-80 °C) were later analyzed for biofilm formation and colony morphology to investigate *P. aeruginosa* adaptation to ETI treatment in a CF airway-like environment.

### Congo red binding assay

The Congo Red binding assay was performed as previously described with modifications [20]. Glycerol stocks were resuspended in 5 mL of Lysogeny broth (LB) and incubated at 37 °C with shaking. Bacterial cultures were then spotted (2 µL) onto Congo Red agar plates (1% tryptone [w/v], 1.5% agar, 40 µg/mL Congo Red, 20 µg/mL Coomassie Brilliant Blue) and incubated at 37 °C for 24 hours. Plates were photographed using Nikon D5300 with Nikon AF-S 60mm f2.8 lens, and colony pigmentation was analyzed using FIJI (ImageJ) with a script for quantification [21]. Briefly, the image is separated into channels, and the pigmentation of whole colonies is measured in the red channel. Data were normalized to the parental PA14 strain using the following ratio: (value of condition – background) / (value of parental strain – background).

### Swarming and swimming motility assay

Swarming and swimming motility assays were performed as described by Ha *et al*. [22,23], respectively, with the following adaptations. For both assays, inoculum was prepared from overnight cultures grown in LB broth at 37 °C with shaking. For swarming motility, supplemented M8 agar plates were prepared by combining autoclaved 5x M8 solution (Na_2_HPO_4_·7H_2_O 64 g/L, KH_2_PO_4_ 15 g/L, NaCl 2.5 g/L) with autoclaved agar at a final concentration of 0.6% (w/v), supplemented with glucose (0.2% w/v), casamino acids (0.5% w/v), and MgSO_4_ (1 mM). Plates (∼25 mL/plate) were poured and allowed to solidify at room temperature for 3-4 hours before use. A volume of 2.5 µL of overnight culture was inoculated at the center of each plate. Plates were incubated upright at 37 °C for 16– 24 hours and photographed for phenotype assessment.

For swimming motility, supplemented M8 agar plates were prepared as described above with a final agar concentration of 0.3% (w/v). Plates were stab-inoculated at the center using a sterile toothpick dipped in overnight culture and incubated upright at 37°C for 16– 24 hours.

### Quantification of SCVs frequency

The proportion of small colony variants (SCVs) was assessed at passage 5 by calculating the ratio of SCVs to the total number of colonies on agar plates. Glycerol stocks from each evolved lineage were recovered, resuspended in 300 µL of LB, diluted, and spread onto LB agar (1.5%) plates to obtain countable colony numbers (30–300 colonies per plate). Plates were incubated at 37 °C for 18 hours, then photographed using the PhenoBooth imaging system.

### Whole genome sequencing

Genomic DNA was extracted from selected samples using the Qiagen DNeasy UltraClean Microbial Kit, according to the manufacturer’s instructions. Whole-genome sequencing was performed using the Oxford Nanopore MinION platform, with sample barcoding carried out using the Rapid Barcoding Kit 96 V14.

The reference genome sequence and annotation files (GFF, GTF, and protein FASTA) for *P. aeruginosa* strain UCBPP-PA14 (ASM1462v1) were retrieved from the NCBI RefSeq database. A custom genomic database was constructed using snpEff v5.2 with the -gtf22 parameter to integrate the GTF annotations with the corresponding reference FASTA sequence. A custom Perl script was used to map gene identifiers to common gene names from the GTF metadata, ensuring consistent nomenclature during variant reporting.

Sequencing data from individual clones isolated at passage 5 were compared to the parental PA14 strain (day 0) to identify mutations acquired during experimental evolution. Variant calling was performed using two complementary pipelines: Medaka (v2.0.1) with the r1041_e82_400bps_sup_variant_v5.0.0_model_pt model and snpEff v5.2. Each sample was processed individually against the PA14 reference genome. Resulting VCF files were processed using bcftools v1.19: each was compressed with bgzip, reheadered to ensure consistent sample identifiers, sorted, and indexed. Functional annotation of the detected variants was performed with snpEff to predict the potential impact of each mutation.

### Quantification of intracellular c-di-GMP using the *cdrA*-lux reporter

Intracellular c-di-GMP levels were quantified using the p*cdrA::lux* transcriptional reporter system, in which the *cdrA* promoter drives expression of the *lux* operon on plasmid pMS402 and is activated by FleQ in response to c-di-GMP. The assay was performed as described by Yaeger *et al* [14], with the following adaptations: individual evolved clones were used as recipient strains and assays were performed in black 96-well plates with clear bottoms (Falcon). Luminescence values were normalized to OD_600_, plotted over time, and AUC was calculated from the resulting curves. The AUC of each clone was compared to that of the parental strain, and results are expressed as log2 ratios.

### Quantification of Pel exopolysaccharide production using the *pel*-GFP reporter

Pel exopolysaccharide production was quantified using a transcriptional reporter system in which the *pel* promoter is fused to *gfp* on a conjugative plasmid. The *pel* promoter was amplified from the previously published pCTX-pro-*pel* plasmid [24] using the FJP_Ppel-GFP_SpeI_For (atccggACTAGTTGCGAGCGGACTGACGGCAAGCAA) and FJP_Ppel- GFP_Xho_Rev (atccggCTCGAGAGCCTACGCGGCAAGGTCGATA) primers. The amplified fragment (∼560 bp) was then cloned in a double restriction enzyme (SpeI, XhoI) digested pSEK-GFP [25] resulting in pSEK-*pel-*GFP. The plasmid was introduced into individual evolved clones by conjugation with the donor strain *E. coli* MFDpir, which requires diaminopimelic acid (DAP) for growth. The donor strain was grown overnight in LB supplemented with DAP (0.3 mM) and tetracycline (10 µg/mL), while recipient clones were grown overnight in LB alone, at 37 °C with shaking. Overnight cultures were mixed at a 1:1 ratio and spotted onto LB agar supplemented with DAP (0.3 mM), then incubated overnight at 37 °C. Spots were scraped, resuspended, and serially diluted before plating onto selective medium containing tetracycline (125 µg/mL) and triclosan (25 µg/mL) to select for transconjugants and counter-select the DAP-auxotrophic donor. For fluorescence measurements, transconjugants were grown overnight in 1 mL of LB supplemented with tetracycline (75 µg/mL) at 37°C with shaking. Cultures were then washed twice with phosphate-buffered saline (PBS) and resuspended in ASM adjusted to OD_600_ of 0.05. Assays were performed in triplicate in black 96-well plates with clear bottoms (Falcon) under static conditions at 37°C using a BioTek Synergy HT plate reader. OD_600_ and GFP fluorescence were recorded every 20 min over 48 hours. GFP fluorescence values were normalized to OD_600_ at each time, plotted over time and AUC was calculated. The AUC of each clone was then divided by the parental AUC and expressed as a log2 ratio.

## Supporting information

**S1 Data**. Variants identified by whole-genome sequencing of individual *Pseudomonas aeruginosa* PA14 clones with distinct morphotypes isolated from DMSO-treated control and ETI-treated lineages.

## Acknowledgments

This work was supported by a Cystic Fibrosis Foundation (CFF) Pilot & Feasibility Award (JEAN21F0) to F.J.-P and a *Programme d’aide au financement* of the Centre de recherche du CHUS (CRCHUS #*93926*) awarded to F.J.-P. and J.-P.C. J.-P.C. holds a Research scholar Junior 2 award from *Fonds de recherche du Québec – Santé* (FRQS). F.J.-P. holds a Research scholar Junior 1 *award* from FRQS (FRQS; #*365777)*. O.G. was supported by grant GONCAL25H0 from the 2025 Student Traineeship Award program of the CFF.

We also thank Dr. Jennifer Bomberger (Dartmouth) for providing ETI used in this study, and Jean-François Lussier (Université de Sherbrooke) for sequencing analyses.

## Notes

### Competing Interest Statement

The authors have declared no competing interest.

